# Selective strolls: fixation and extinction in diploids are slower for weakly selected mutations than for neutral ones

**DOI:** 10.1101/016881

**Authors:** Fabrizio Mafessoni, Michael Lachmann

## Abstract

In finite populations, an allele disappears or reaches fixation due to two main forces, selection and drift. Selection is generally thought to accelerate the process: a selected mutation will reach fixation faster than a neutral one, and a disadvantageous one will quickly disappear from the population. We show that even in simple diploid populations, this is often not true. Dominance and recessivity unexpectedly slow down the evolutionary process for weakly selected alleles. In particular, slightly advantageous dominant and mildly deleterious recessive mutations reach fixation more slowly than neutral ones. This phenomenon determines genetic signatures opposite to those expected under strong selection, such as increased instead of decreased genetic diversity around the selected site. Furthermore, we characterize a new phenomenon: mildly deleterious recessive alleles, thought to represent the vast majority of newly arising mutations, survive in a population longer than neutral ones, before getting lost. Hence, natural selection is less effective than previously thought in getting rid rapidly of slightly negative mutations, contributing their observed persistence in present populations. Consequently, low frequency slightly deleterious mutations are on average older than neutral ones.

**Author Summary:** A common assumption among geneticists is that neutral alleles survive longer in a population than selected variants: negative selection would rapidly lead to the extinction of deleterious mutations, while advantageous alleles under positive selection will spread in the population till fixation. Here we show that unless an allele is perfectly codominant, these assumptions are often incorrect. Under weak selection, even in the simplest models, incomplete dominance and recessivity are sufficient to determine slower fixation and extinction times than for a neutral allele. These seemingly paradoxical results suggest that mildly deleterious mutations accumulate at the population level, and that nearly neutral mutations behave very differently from strongly selected ones. Furthermore, a fraction of selected regions would show opposite patterns for many standard statistics used to detect genomic signatures of positive selection, remaining virtually impossible to detect.

## Introduction

A new allele emerging in a finite population usually has two possible fates – extinction or fixation. Selection affects the probability with which these occur, and how long it will take. Thus, an advantageous allele has an increased chance to fix, due to positive selection, while a deleterious mutations has an increased chance of extinction. In both cases, the time till fixation and extinction is commonly thought to decrease with the strength of selection[1]. Kimura [1, 2] and Ewens [3] applied diffusion theory to model finite populations, obtaining keystone approximations for the neutral case, in absence of selection, or when selection is fairly strong. Recently it has been pointed out that in haploid models, in presence of frequency-dependent fitness, the time to fixation of a positively selected allele can *increase* with the strength of selection [4, 5]. In diploids, a newly arising mutation is usually expected to have a time to fixation longer than a neutral one in case of overdominance, when the heterozygote has an higher fitness than the two homozygotes – a case usually referred to as heterozygote advantage or more generally balancing selection [6]. In case of overdominance, both alleles would be maintained in an infinite population. In this paper we unify these results, showing that in diploids, certain classes of mutations behave as *slow sweeps*: the time to fixation of a positively selected allele *A* can be longer than in the neutral case, provided that selection is weak and only requiring incomplete dominance (i.e. the heterozygote *AB* has higher fitness than the disfavoured homozygote *BB* but lower or equal to the favored homozygote *AA*).

Maruyama [7] has already shown that the time to fixation for deleterious alleles decreases as selection becomes more negative. For a simple haploid model with a selected allele of selection strength 1 + *s* vs. the wild type with fitness 1, the time to fixation is equal for *s* = +|*x*| and *s* = −|*x*|. For diploids the same phenomenon occurs, provided that the dominance relationship between alleles is reversed [3]. Hence, our findings extend to the case of mildly deleterious and recessive mutations.

We also investigate the trajectories conditional to extinction of newly arising deleterious mutations. Deleterious mutations are thought to be rapidly purged from a population. Combining our results, we demonstrate that mutations with these features, likely constituting a large fraction of newly arising ones, not only have longer fixation times, but survive in a population longer than neutral ones. Hence we show that negative selection is less rapid than usually thought in getting rid of midly disadvantageous alleles, and that these are on average older than neutral ones.

When a selected mutation spreads in a population till fixation, surrounding sites are carried along, a phenomenon called genetic hitchhiking. Since a selected mutations usually spread rapidly and less time is available for recombination, the genetic diversity around a positively selected site is commonly expected to decrease, a phenomenon called *selective sweep*[8, 9]. Most statistics used to detect genetic signatures of positive selection rely on this assumption [10, 11, 12, 13]. We show that these signatures are reversed in *slow sweeps*, with increased genetic diversity compared to neutrally fixing allele. We then explore the question of how often such a slowdown would occur in a population, by using different assumptions about the distribution of selection and dominance effects in nature.

## Results

### Dominant weakly advantageous alleles reach fixation more slowly than neutral ones

We investigate a classic single locus two-allele Wright-Fisher model. The wildtype homozygote with genotype *aa* has fitness 1, and we study the fate of a newly introduced allele *A*. The fitness of the homozygote *AA* is 1 + *s* where *s* denotes the selection coefficient, while the heterozygote *Aa* has fitness 1 + *hs*, where *h* denotes dominance. In the absence of mutations, extinction and fixation, when the frequency of allele *A* equals 0 or 1, respectively, are two absorbing states. We explored the conditional expected time till either one or the other event occur, with diffusion approximation and simulations, referring to them for brevity as extinction and fixation time, respectively. In Fig.1 we show diffusion approximations for the fixation time of weakly selected positive (*s >* 0) and negative (*s <* 0) mutations, relative to neutrality. For completely dominant positive (*h* = 1) allele *A* a peak in the time of fixation is observed around *N*_*e*_*s* ≃ 1, where *N*_*e*_ is the variance effective population size. An analogous effect is observed for completely recessive (*h* = 0) deleterious mutations. This result depends only on *N*_*e*_ *s* and *h*, hence it holds also for small *N*_*e*_ (see Fig.S7 for simulations). Remarkably these effects occur even for intermediate levels of dominance, although to a smaller degree as *h* approaches 1/2. The symmetry of fixation times has been already noticed by Maruyama [7], who showed that inverting the coefficient and the sign of the selection coefficient keep unchanged the fixation time. However this symmetry is centered at *s* = 0 only when *h* = 0.5. Hence a balancing selection-like pattern, with increased fixation times compared to neutrality, is observed in a non-trivial region in the *s − h* parameter space (Fig. 1): increased fixation times occur in a hourglass-shaped parameter region, rather than in a simpler rectangular fashion (overdominance, *h* > 1, *s* > 0), due to the stochastic slowdown shown here for weak selection, and the decrease in fixation time for overdominant strong selection described by Robertson [14].

**Figure. 1:**
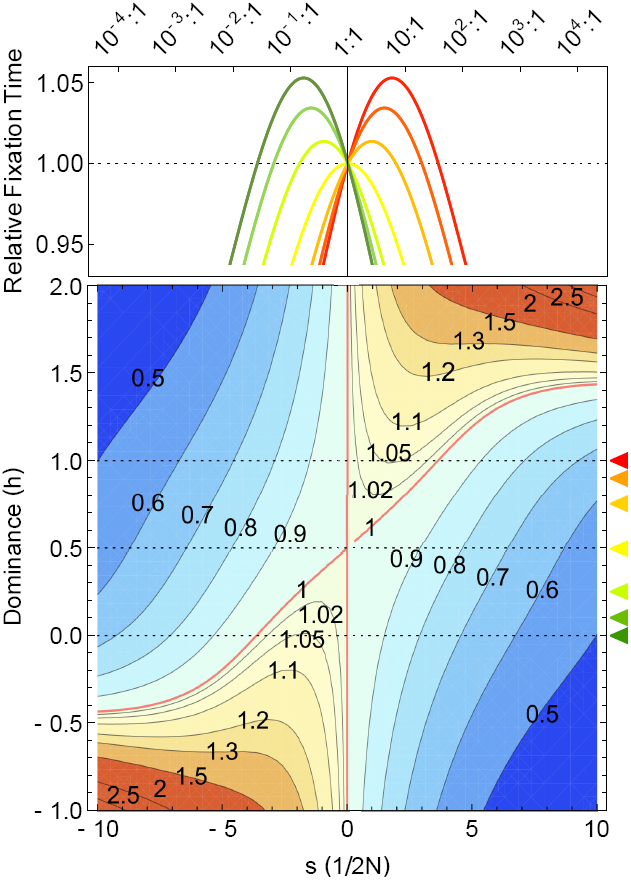
**Time of fixation relative to neutrality for different selection coefficients (x-axis) and dominance for general a population of size** *N*_*e*_. a) In the top panel different curves indicate different levels of dominance, from the red to the green, *h* is equal to 1, 0.9, 0.75, 0.5, 0.25, 0.1 and 0, as indicated by the colored triangles in (b). On the top y-axis selection is also measured in terms of the odds for a site to be fixed in *A* rather than *B*, at the stationary state, (for *h* = 0.5 and 2*N*_*e*_ = 10^4^). b) Time of fixation relative to neutrality for different values of positive selection (x-axis) and dominance (y-axis). A red contour (fixation times equal to neutrality) separates shorter (shades of blue) from longer (warm shades) fixation time than under neutrality. Dotted lines separate (from top to bottom) overdominance, dominance, recessivity and underdominance.

### An intuitive explanation of the stochastic slowdown in diploids

Why does this counter-intuitive phenomenon occur? Altrock et al. [5], pointed out that in haploids the sojourn times, the amount of time spent at the different population frequencies, increase at higher frequencies. Here we provide a simple explanation for the stochastic slowdown in diploids, by looking at the conditional transition probabilities. A conditional trajectory is slower whenever the frequency of the selected allele decreases, only to increase again later and eventually reach fixation. Therefore the fixation probability of a mutation going to lower frequencies has a key role in the fixation time: in the case of positive selection dominance increases the probability that once a drop in frequency occurs the selected allele *A* will reach fixation anyway. This occurs since even if there is a higher chance for *A* to be paired with the less advantageous allele *a*, the fixation probability is still relatively high because the average fitness of the heterozygote is biased toward the more advantageous allele. This effect is apparent only for weak selection, since when *s* is strong, the probability to decrease in frequency, compared to the probability of increasing, it is so low that this effect can be neglected. A similar explanation is applied to recessive deleterious mutations. In this case recessivity masks the deleteriousness of the selected allele *A*, so that when a drop in frequency occurs, its absolute fitness will increase since it will be biased towards the advantageous allele. Hence, its probability of fixation would be relatively high [15, 16].

In order to illustrate this intuitive explanation, we use a simplified Moran process, with fitness mirroring that of a diploid biallelic population with 2 *Ne* alleles [17]. In a Moran process the number of *A* alleles, indicated as *i*, can change at most one allele at the time, so that the only non-zero transition probabilities are those of moving from *i* to *i* + 1, from *i* to *i* − 1, or of remaining in *i*. In our simplified process, we ignore steps that do not change the frequency of the allele. We denote the fixation and extinction probabilities in *i* = 0 and *i* = 2*N*_*e*_ as 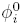 and 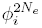, respectively, and the transtion probabilities of a decrease or an increase in frequency as 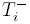 and 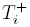. The conditional transition probabilities, given that fixation will eventually occur, are obtained by multiplying the unconditional transition probabilities by the probability of fixation once in the new state, and by normalizing for the fixation probability over all the possible destination states. In the simplified Moran process, the conditional probabilities of a decrease or an increase in the number of *A* alleles given fixation, are simply equal to the areas of the rectangles 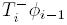, and 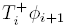 respectively, normalized by their sum (Fig.2, 3). Hence for each *i* and given combination of *s* and *h*, a slowdown occurs whenever the probability of a decrease in frequency conditional on fixation is larger than under neutrality (*s* = 0) i.e. the left rectangle 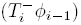 is larger than the right one 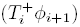, if compared to neutrality. This condition can be simply expressed as 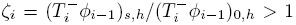 where 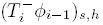 is calculated for selection and dominance coefficients *s* and *h*. In figure 2 we illustrate what happens in the case of positive selection, with the neutral case represented for comparisons as semi-transparents rectangles and arrows in all figures, and in blue in Fig.2b. It is easy to observe how dominance buffers the decrease in frequency by increasing the probability of fixation for lower *A* frequencies, thus increasing the height of the rectangle 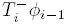 (Fig.2a). When selection is stronger, the transition probability 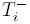 is much smaller, and the area of the rectangle 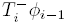 cannot be compensated by increasing its relative height, compared to neutrality (Fig.2.d). Recessivity exerts the opposite effect, accelerating fixation (Fig.2c). Vice-versa, recessivity determines longer fixation times in the case of a deleterious *A* allele (Fig.3). The stochastic slowdown is stronger at higher frequencies (Fig.S3-S4),as it can be also seen from the distribution of conditional sojourn times reported for the Wright-Fisher model (Fig.S1).

**Figure. 2:**
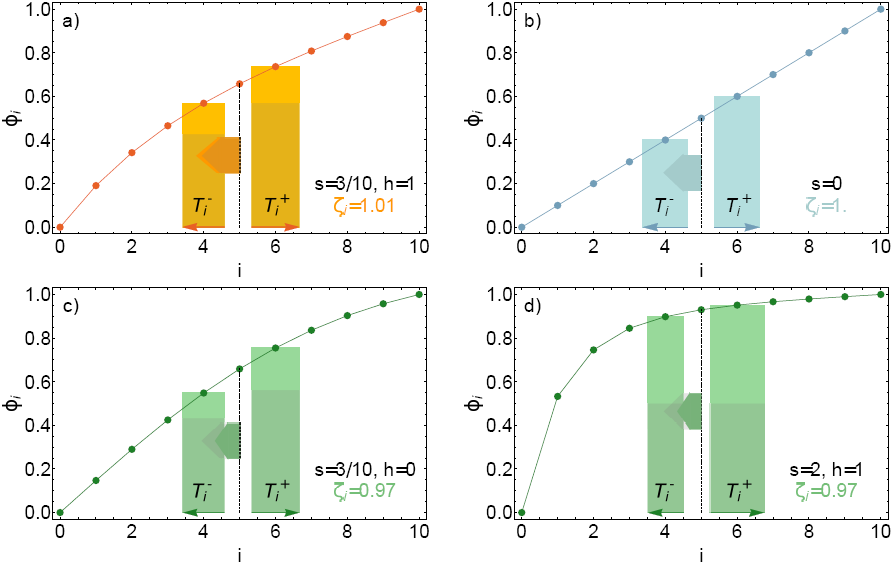
**Slowdown for a simplified Moran process with** *i* = 5 *A* **selected alleles for positive selection.** The number of *A* allele is represented on the x-axis, the fixation probability for the different states on the y-axis. The coloured rectangles indicate the not-normalized transition probabilities of an increase or a decrease in frequency conditional to fixation, given by product of the transition probability (thin arrows) and the fixation probability for the arrival state. The length of the thick arrow is proportional to (*ζ_i_*)^10^. Transparent arrows and rectangles indicate the same quantities for the neutral case. Figures a-d represent respectively a weak dominant selection case, neutrality, weak recessive and strong selection.

**Figure. 3:**
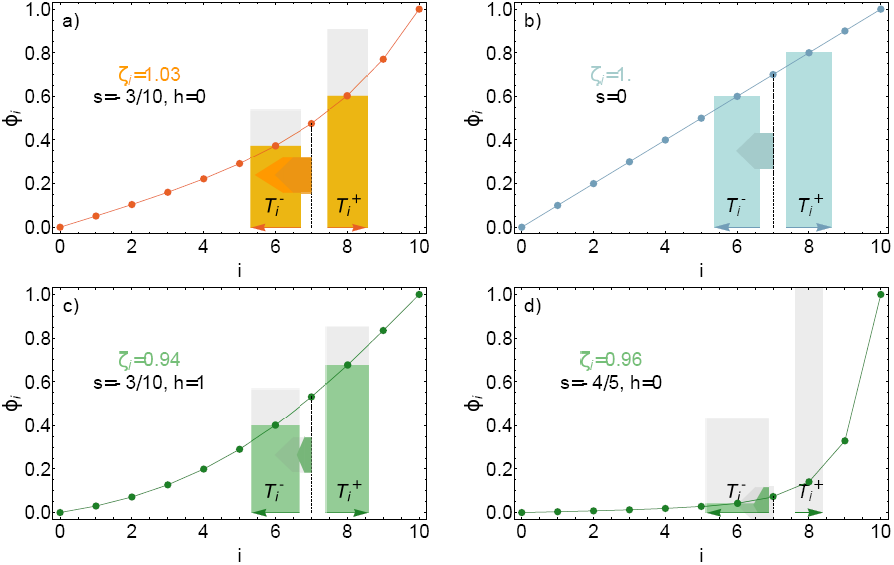
**Slowdown for a simplified Moran process with** *i* = 5 *A* **selected alleles for negative selection.** Figures a-d represent respectively a weak recessive selection case, neutrality, weak dominant and strong selection. Figure details are the same as in Fig.2

#### *Slow sweeps* signatures

Consistenly with the longer time till fixation, genetic diversity is increased compared to neutrality around a selected site, in opposite fashion to classical selective sweeps (Fig.4). Hence, even classical statistics to detect positive selection as Tajima’s D show opposite pattern for slow and classical selective sweeps. We used hypothetical distribution of dominance and selective effects to estimate the fraction of fixation events due to either slow or classical selective sweeps (Fig.S5). We assumed the distribution of selective effects (DFE) estimate in Racimo [18], and explored a truncated normal distribution for dominance effects. When the variance of the latter is high, this converges on a uniformous distribution, and the fraction of fixation events due to slow sweeps is about one half (Fig.S5.e). This fraction decreases rapidly as dominant mutations are rarer. Since the DFE is estimated for mostly deleterious mutations, we also conservatively take into consideration the possibility that the distribution of selection coefficients is much more skewed towards stronger effects for positive selection. Hence we considered exponentially DFE with mean five and ten times higher than in Racimo et al.[10]. In these cases the fraction of fixation events due to slow sweeps is lower, when *h* is uniformly distributed about 1/5 and 1/10, respectively (S5.f).

**Figure. 4:**
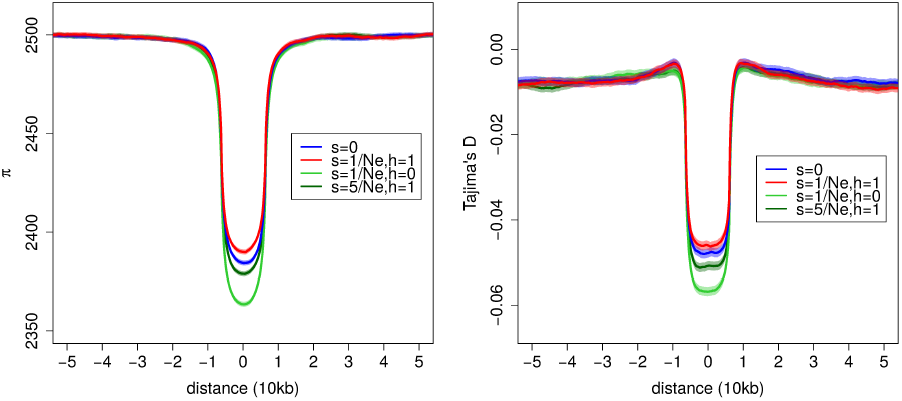
**Patterns of nucleotide diversity** (*π*) **and Tajima’s D around the site of a selected allele that just reached fixation.**We considered a neutral (blue), dominant (red) and recessive (dark green) weakly selected allele, and a dominant allele with stronger selection (light green) as indicated in the legend. The population size is *N*_*e*_ = 10000, mutation rate 1.2 * 10^−8^. Shaded regions indicate the 95% confidence interval of 5 * 10^5^ simulations.

### Recessive weakly deleterious alleles persist in a population longer than neutral ones

It is estimated that most newly arising mutations are slightly deleterious and recessive, consistenly with the nearly neutral theory of molecular evolution[16, 19, 18, 20, 21]. A remarkable implication of the selective slowdown is that these mutations disappear more slowly than neutral ones from a population: the fixation of a slightly advantageous selected mutation is equivalent to the extinction of a slightly deleterious recessive one. Furthermore we have seen that the slowdown process is stronger for high frequency of the advantageous mutation, hence for low frequencies of the deleterious one. Hence we can ask what is the time till extinction of a newly appeared mutation. This is also determined by the arrival time, the time that this mutation takes to initially invade. We have already observed how this process is slower, when considered till fixation. Therefore we hypothesize that the extinction time of slightly deleterious recessive mutation is longer than neutrality. We show this seemingly paradoxical result in figure 5 for *Ne* equal to 10^4^. The extinction times are longer than neutrality for recessive deleterious mutations. When recessivity is complete, even mutations with a selective coefficient almost as deleterious as −10/2*N*_*e*_ disappear more slowly than neutral ones. While conditional fixation times relative to neutrality are dependent only on *h* and *N*_*e*_ *s*, the stochastic slowdown for extinction is stronger for smaller absolutte population sizes. For population with 2*N*_*e*_ equal to 100 the extinction time can be even 10% longer than neutrality, while for populations with *N*_*e*_ larger than 10^4^ the effect of the stochastic slowdown is at most 5%. Consistenly with this phenomenon, recessive deleterious mutations have longer unconditional sojourn times than neutral ones at low frequencies (Fig.6a-b). Thus, contrarily to what is generally thought, these alleles are on average older than neutral ones (Fig.6c-d). This phenomenon is stronger at low frequencies, where the range of selection coefficients for which deleterious alleles are older expands (Fig.6c).

**Figure. 5:**
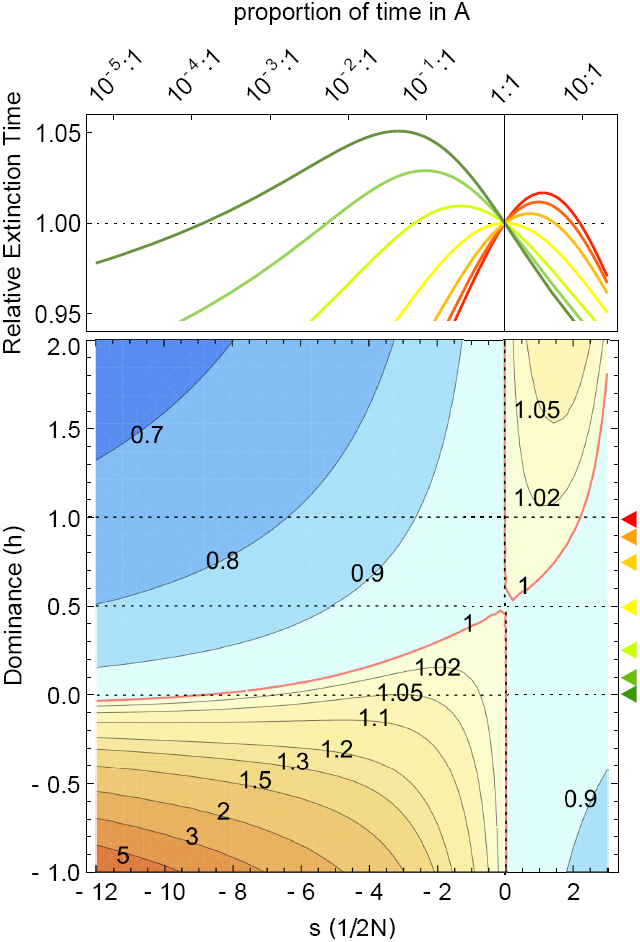
**a) Time of extinction relative to neutrality for different selection coefficients (x-axis) and dominance for a population of size** 10^4^. In the top panel different curves indicate different levels of dominance, from the red to the green, *h* is equal to 1, 0.9, 0.75, 0.5, 0.25, 0.1 and 0, as indicated by the colored triangles in (b). On the top y-axis selection is also measured in terms of the odds for a site to be fixed in *A* rather than *B*, at the stationary state, (for *h* = 0.5 and 2*N*_*e*_ = 10^3^). b) Time of fixation relative to neutrality for different values of positive selection (x-axis) and dominance (y-axis). A red contour (fixation times equal to neutrality) separates shorter (shades of blue) from longer (warm shades) fixation time than under neutrality. Dotted lines separate (from top to bottom) overdominance, dominance, recessivity and underdominance.

**Figure. 6:**
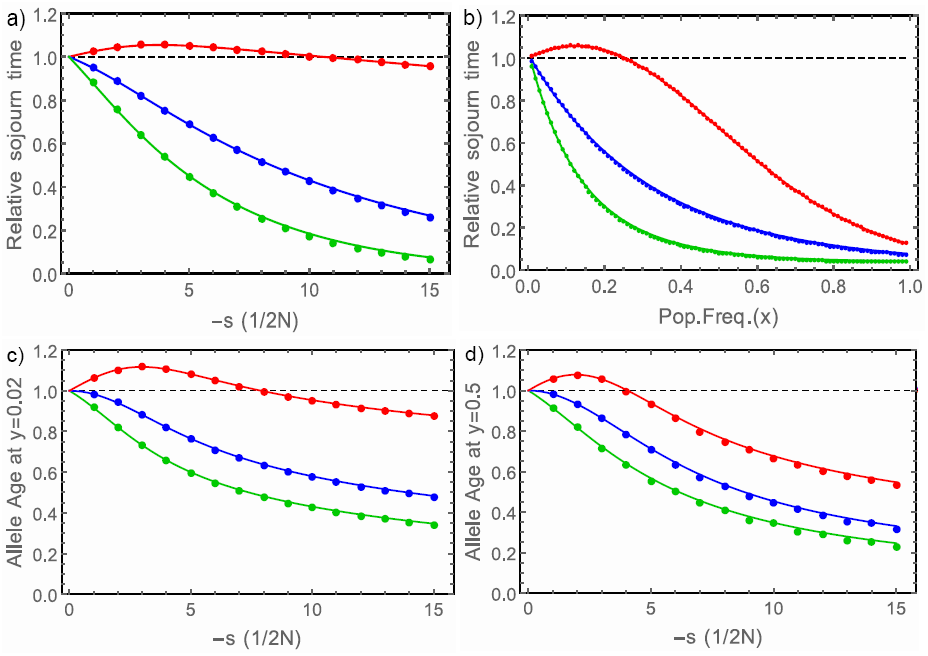
**Sojourn times (a-b) and ages (c-d) of a mutated allele, relative to neutrality, for completely recessive (red), codominant (blue) and dominant (green) deleterious mutation.** Dots indicate the average of 10^6^ simulations, while lines diffusion approximations. In (a,c,d) different selective coefficient are shown on the *x*-axis, while for (*b*) the relative sojourn times are shown for different population frequencies, with *s* = −2/*N*. In Fig (c) and (d) the population frequency is respectively 0.02 and 0.5.

## Discussion

Our results show that the mild frequency-dependence of the fitness of a selected allele, even when only due to small levels of recessivity and dominance, is enough to alter the expected time till fixation and extinction of nearly neutral mutations. A common assumptions in population genetics is that positive selection leads to a shortening of fixation times [22]. Most methods aimed at detecting molecular signatures of positive selection rely on this assumption [9, 23]. Here we showed that weakly selected dominant alleles, with selection coefficients
around 2/2*Ne*, violate the assumption of a rapid spread in the population. These alleles behave as *slow sweeps*, on average reaching fixation more slowly than neutral ones. This is coherent with the difficulties in detecting
hard sweeps, usually explained in terms of polygenic adaptation [24]. Despite being well known that selective sweep can be hard to find when selection is weak, here we show that the expected signatures of these slow selective events are not only weak, but even in the opposite direction of what expected: diversity around a fixed selected allele is higher than around a fixed neutral one. Identifying fixation events subjected to a stochastic slowdown is unlikely, due to its limited effects (about 5%). However, our result rather show that selective events involving certain classes of mutations are inherently elusive and virtually impossible to detect with standard methods.

What is the fraction of such fixation events? First of all we observed how this phenomenon appear for mutations under weak selection (*N*_*e*_ *s* ≃ 1). Hence, for very large populations this phenomenon might be effective only for alleles with very weak effects. The fixation probability of weakly selected mutations is relatively low, only 3 times higher than neutral ones for *N*_*e*_ *s* = 1 and *h* = 1, although at the stationary-state each site spend most time in the selected state (85% of the time for the same mutation). Despite the relatively small fixation probability, the distribution of selection coefficients of newly arising mutations is likely exponential-like, with the vast majority of mutations having weak effects [18, 19]. For this reason, slow sweeps might constitute a substantial fraction of fixation events. An exact estimate of the fraction of selective events due to slow sweeps is limited by the lack of a precise knowledge of the distribution of dominance effects for newly arising mutations. However, exploring hypothetical distributions, we estimate that the fraction of *slow sweeps* should vary between one half or one tenth, with conservative assumptions.

We have also shown that weakly deleterious recessive alleles have longer average extinction times than neutral ones, both in the case of common deleterious variants and for newly arising mutations. Although there is not a clear picture of the effects of dominance for positively selected mutations, most deleterious ones are thought to be recessive [21]. For this reason, an increase in the length of extinction times of such mutated alleles is extremely relevant, implying that purifying selection is less rapid than previously thought in removing disadvantageous variants. For this class of mutations the sojourn times, and thus in turn the site-frequency-spectra, are affected. In particular the sojourn times at low frequencies are longer compared to neutrality, implying an accumulation of weakly deleterious variants at the population level[25]. Nevertheless, empirical studies failed to detect our theoretical predictions of older weakly recessive deleterious mutations compared to neutral ones [26, 27, 28]. This is probably due to the difficulties in determining exactly which mutations are slightly deleterious, and which are recessive versus codominant dominant. Future studies, and better estimates of the distribution of selective and dominance effects, might be able to confirm our theoretical predictions.

Hence our study shows qualitative patterns indicating that the speed of the stochastic dynamics of selected alleles is often affected in a non trivial way by dominance and recessivity. These results suggest new testable predictions about the permanence of deleterious mutations in current populations, and challenge current methods to detect signatures of positive selection in genomes.

## Supporting Information

### S1 Text

#### Materials and Methods

**Diffusion Approximation** We assume a Wright-Fisher population in the limit of large population size and use Diffusion theory as described in Ewens (2004). The density function *ϕ*(*t, p*) of the time *t* until absorption, starting from frequency *p*, satisfies the backward Kolmogorov equation:

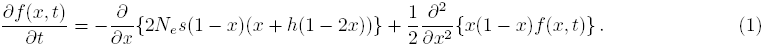

We express the unconditional sojourn time between *a* and *b* as 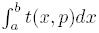. When *a* and *b* corresponds to the absorbing states 0 and 1, we obtain the unconditional time till absorption:

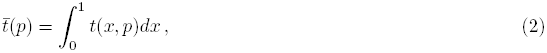

where

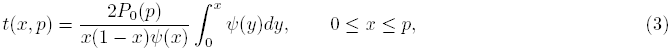

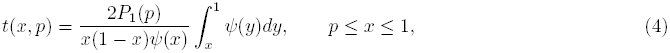

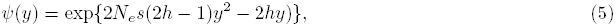

and the probability of fixation *P*_1_(*x*) can be expressed as:

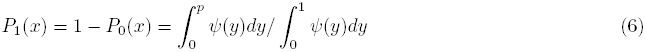

The mean conditional time till fixation or extinction can be directly obtained by conditioning sojourn times to fixation or extinction, respectively, and then integrating over all population frequencies:

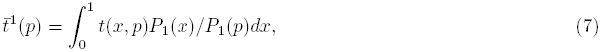

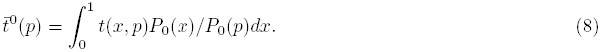

Following Maruyama and Kimura, the age of a mutant allele can be calculated by integrating over the density of sojourn times at *z*, given that the current frequency is *x*:

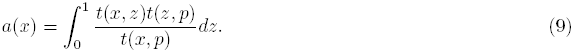

**Moran process** We model a birth-death Moran process as in Altrock et al.2012, for which fixation and extinction times can be calculated analytically. We consider a bi-allelic haploid well-mixed populazion of size 2*N*. We denote the number of *A* alleles as *i* and *B* alleles as 2*N − i*. In order to mimick the fitness structure of a diploid population we consider the birth and death probability of birth of type *A* respectively equal to:

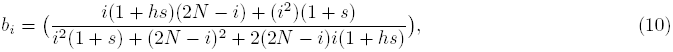

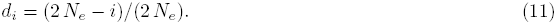

Hence the transition probabilities to go from *i* to *i* + 1, from *i* to *i -* 1, and to stay in *i* are equal to:

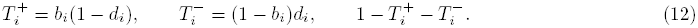

We also consider the simplified Moran process introduced in the main text. In this case the focal allele is not in an absorbing state, its frequency is never the same in two consecutive time steps. Thus if at time *t* there are *i A* alleles, with *i* ≠ 0 and *i* ≠ 2*N*, at time *t* + 1 there can be only *i* − 1 or *i* + 1 *A* alleles. Hence we normalize the new transition probabilities as 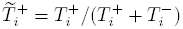 and 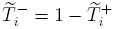.

Keeping 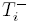 and 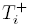 to describe in general unconditional transition probabilities, for both the general and the simplified model, the arrival probability at state *j* can be obtained by solving the recursion equation (Ewens, 2004):

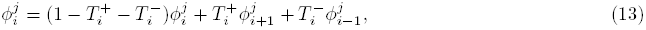

thus:

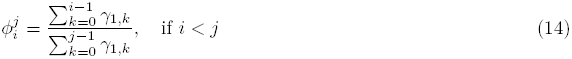

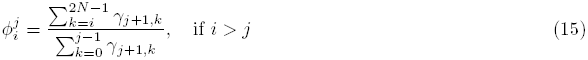

where 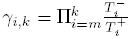. The fixation and extinction probability are then denoted as 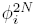 and 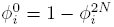.

The conditional fixation and extinction time, 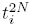 and 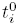 can be calculated similarly to the sojourn times, as in Altrock et al.2012 [5]. The average sojourn time in *j*, starting in *i*:

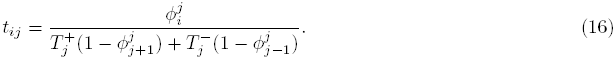

We notice that for a transition probability *T*_*ij*_ from state *i* to state *j*, the conditional transition probabilities, given that the absorbing state *a* is reached, can be obtained as 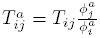. Hence the conditional fixation 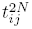 and extinction 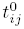 sojourn times can be obtained by conditioning as in Ewens 2004:

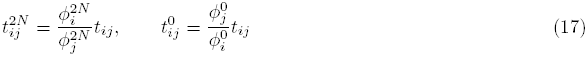

The average time till absorption, fixation and extinction are simply the sum of the unconditional, conditioned on fixation and on extinction sojourn times in each state:

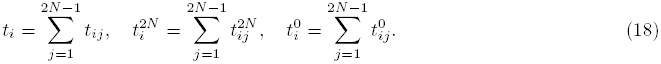

**Simulations** We used SLiM[29] and custom code for unconditional simulations, and MSMS [30] for simulations conditioned on fixation. Wright-Fisher simulations have been performed for either 10000 or 50 individuals. For unconditional simulations (sojourn times, age of segregating alleles) we performed 11^7^ independent simulations. Out of these simulations we subsampled 10^6^ runs for fixation and 10^7^ runs for extinction to verify the accuracy of diffusion approximations.

**Figure. S1.**
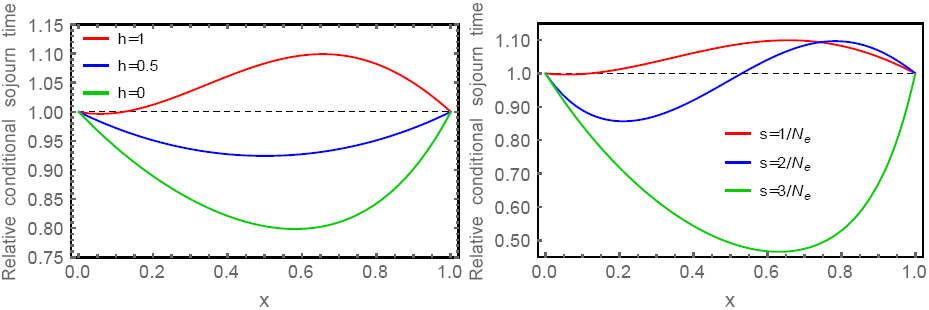
**Diffusion approximation of the conditional sojourn time relative to neutrality at different population frequencies (x).** On the left side the sojourn times are shown for *s* = 1*/N_e_*, for a single emerging allele in the case of complete dominance (red), codominance (blue) and recessivity (green). On the right side the sojourn times are shown for a completely dominant allele with different values of *s*. In the case of weak selection and *h* > 0.5 the conditional sojourn times are longer than under neutrality, especially at higher population frequencies.

**Figure. S2.**
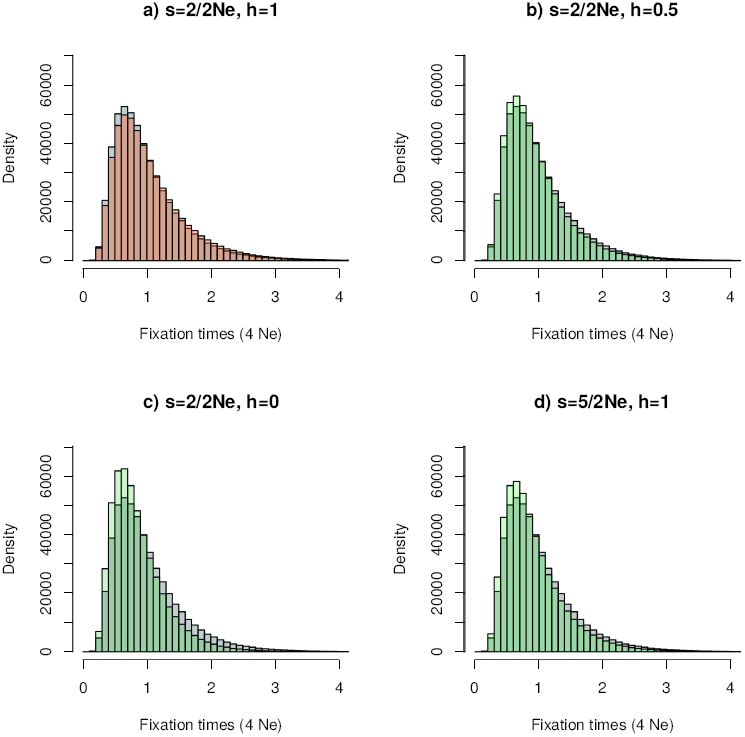
**Distribution of fixation times for** 5 * 10^5^ **Wright-Fisher simulations with** *N* = *N*_*e*_ = 10^5^ **and parameters as indicated in the figures.** Neutral distributions are indicated in gray, while red and green indicate combinations of *s* and *h* for which an increase (a) or a decrease in conditional fixation time is expected, respectively (b-d).

**Figure. S3.**
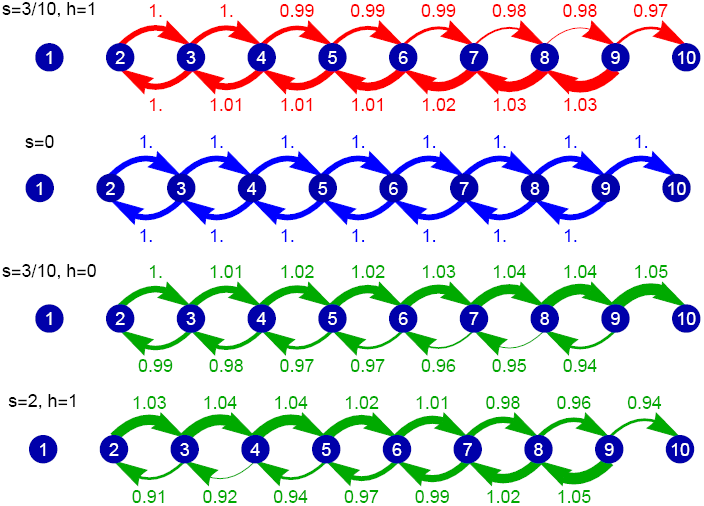
**Transition probabilities conditional on fixation of an increase (top arrows) or a decrease (bottom arrows) in** *A* **alleles in the simplified Moran process** with 2*N* = 10 for values of *h* and positive selection corresponding to Fig. 2

**Figure. S4.**
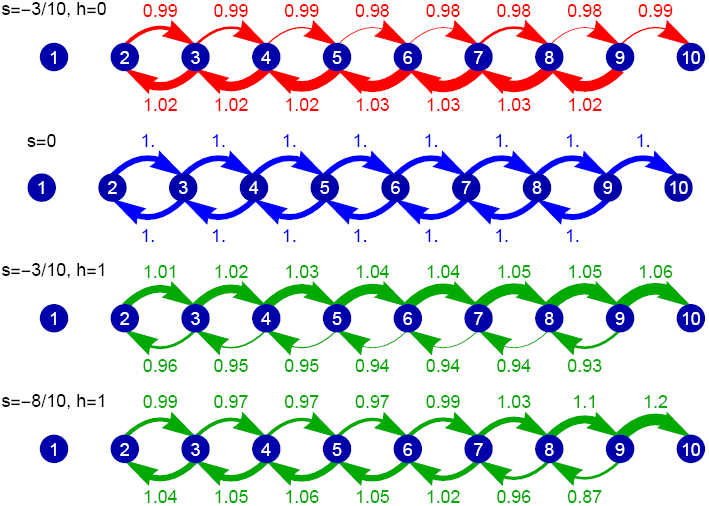
**Transition probabilities conditional on fixation of an increase (top arrows) or a decrease (bottom arrows) in** *A* **alleles in the simplified Moran** process with 2*N* = 10 for different values of *h* and negative selection corresponding to Fig. 3

**Figure. S5.**
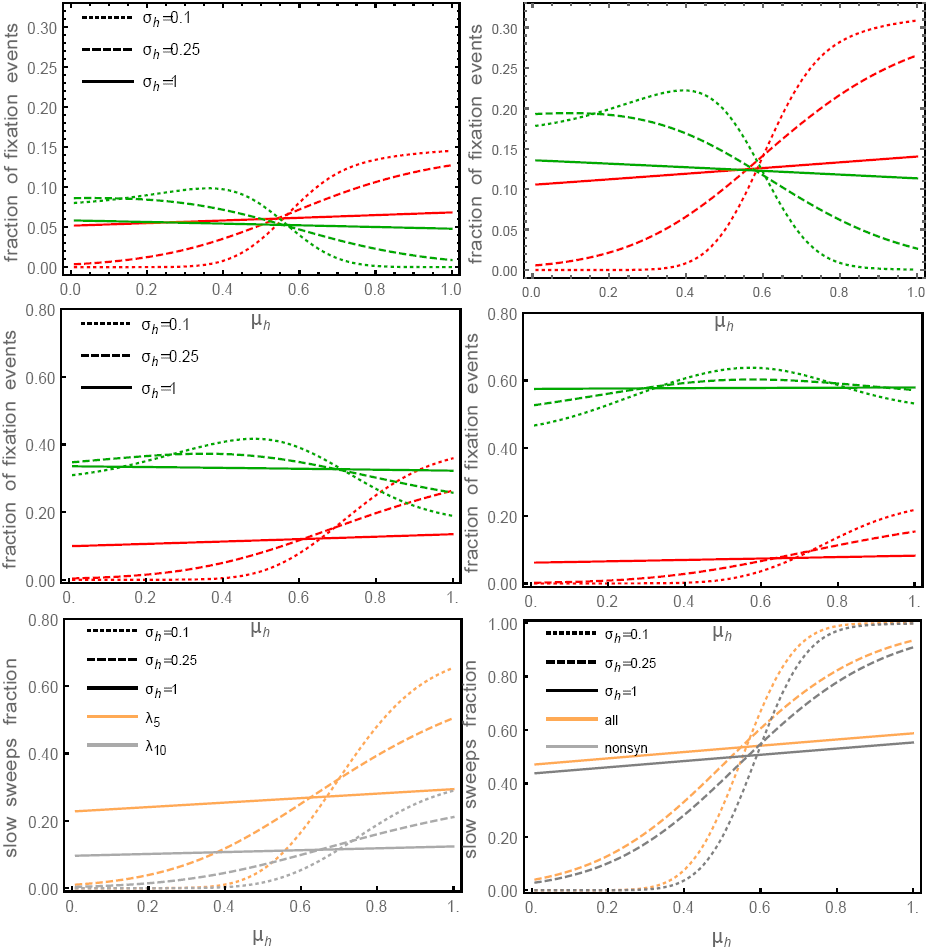
**Fraction of fixation events (y-axis) due to drift and selection with longer (red) and shorter (green) fixation times than neutrality for different distribution of selective effects (DFE) and dominance.** We exclude overdominance and underdominance cases, assuming dominance to be distributed following a truncated normal distribution with mean *μ_h_* and variance 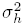. Selection follows the DFE estimated in Racimo et al.[10] at the genome wide level, for all mutations (a), and non-synonymous mutations (b). We also considered exponentially distributed selection coefficients, with mean 5 (c) and 10 (d) higher than estimated in Racimo et al.[10]. For each value of *s* and *h* we calculated the fraction of neutral fixation events and selective ones as 1/2*N*_*e*_ and *P*_1_(1/2*N*_*e*_) − 1/2*N*_*e*_, respectively.

**Figure. S6.**
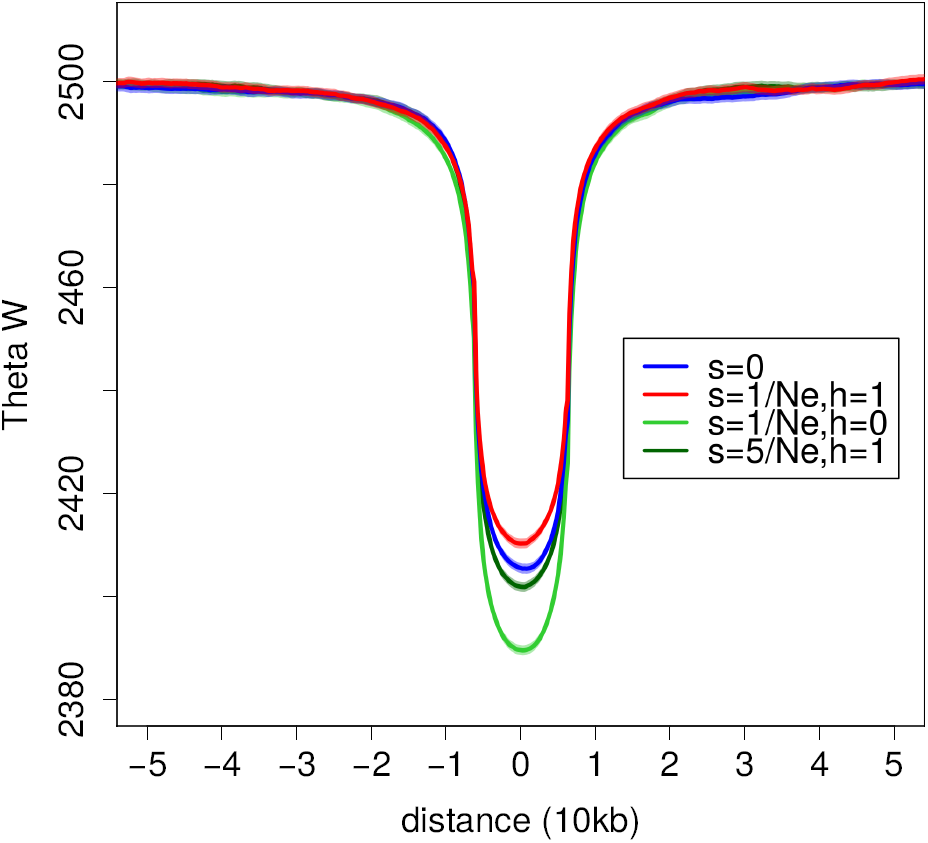
**Watterson’s** *θ* **around the site of a selected allele that just reached fixation.** We considered a neutral (blue), dominant (red) and recessive (dark green) weakly selected allele, and a dominant allele with stronger selection (light green) as indicated in the legend. Shaded regions indicate the 95% confidence interval of 5 *\** 10^5^ simulations for a population with *N*_*e*_ = 10000, mutation rate 1.2 * 10 ^−8^.

**Figure. S7.**
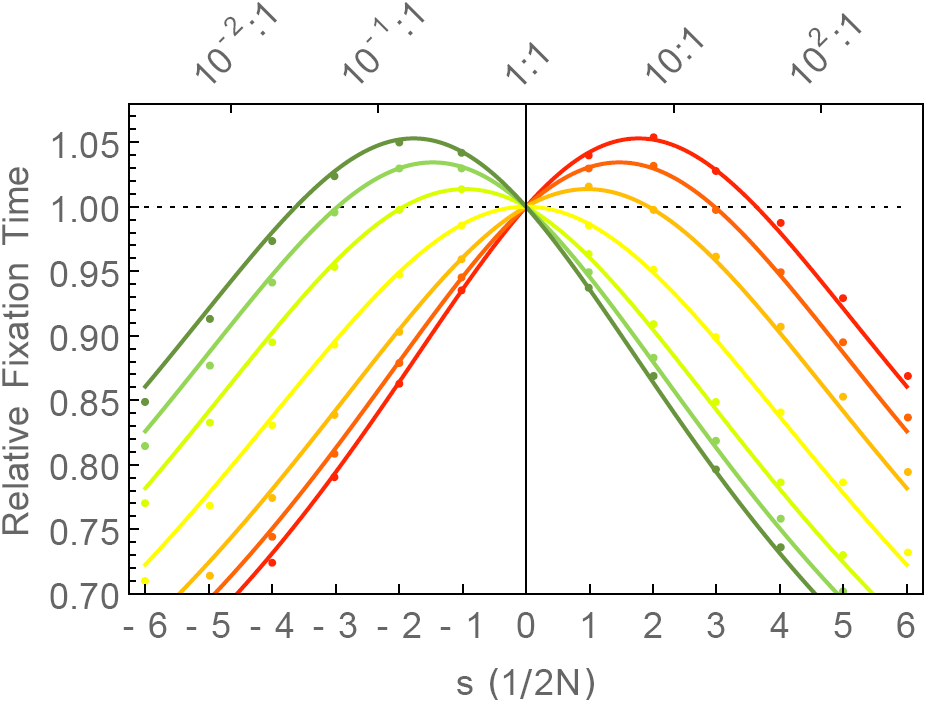
**Diffusion approximation (continous lines) and the average of** 10^5^ **simulations (dots) for the fixation times relative to neutrality for a single initial allele in a population with** *Ne* = 50. Error bars indicate the 95% confidence interval. Colors indicate value of *h* corresponding to figure 1.

**Figure. S8.**
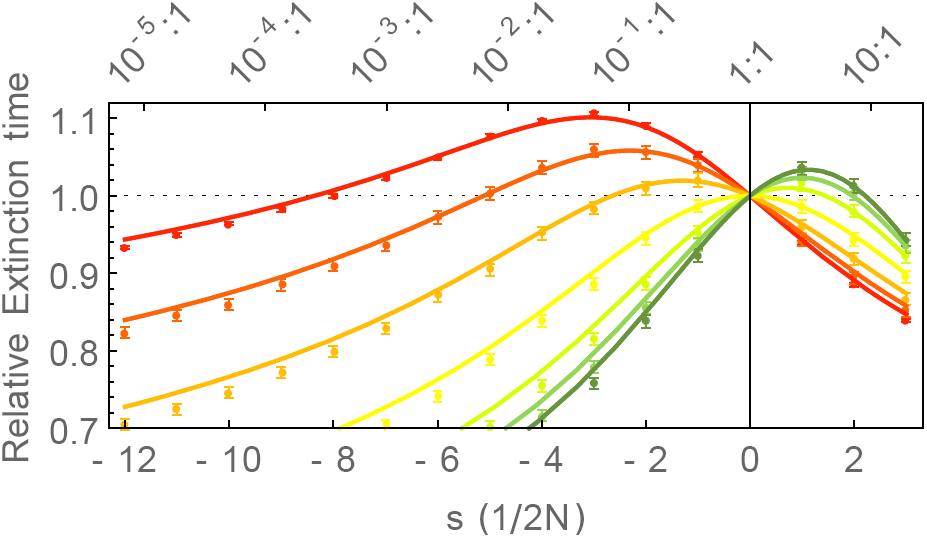
**Diffusion approximation (continous lines) and the average of** 10^7^ **simulations (dots) for the extinction times relative to neutrality for a single initial allele in a population with** *Ne* = 50. Error bars indicate the 95% confidence interval. Colors indicate value of *h* corresponding to figure 1.

